# Hepatitis C Virus Regulates its Replication by Maturing miR-122 Through Akt-Dependent Phosphorylation of KSRP

**DOI:** 10.1101/654392

**Authors:** Camille Baudesson, Céline Amadori, Hassan Danso, Flora Donati, Quentin Nevers, Tony Durand, Matthieu Lemasson, Arielle R. Rosenberg, Michele Trabucchi, Abdelhakim Ahmed-Belkacem, Patrice Bruscella, Jean-Michel Pawlotsky, Cyrille Féray

## Abstract

The liver-specific micro-RNA-122 (miR-122) is required for the replication of hepatitis C virus (HCV). The direct interaction between miR-122 and the 5’ untranslated region of the HCV genome promotes viral replication and protects HCV RNA from degradation. Because HCV RNA is its own substrate for replication, infected cells are submitted to the sequestration of increasing levels of miR-122 and to global de-repression of host miR-122 mRNA targets. Whether and how HCV regulates miR-122 maturation to create an environment favorable to its replication remains unexplored. We discovered that Akt-dependent phosphorylation of KSRP host protein at Serine residue 193 is essential for miR-122 maturation in hepatocytes. Moreover, we showed the existence of a reciprocal regulation loop where HCV replication can modulate the proviral effect mediated by KSRP-dependent maturation of miR-122. These data support a mechanism by which HCV regulates the expression of miR-122 by hijacking KSRP, thereby fueling its own replication.

## Introduction

Hepatitis C virus (HCV) is a positive-strand RNA virus causing chronic liver disease, including chronic hepatitis, cirrhosis and/or hepatocellular carcinoma (Hoofnagle, 2002). The HCV RNA genome is uncapped, non-polyadenylated, and contains highly structured 5’- and 3’-untranslated regions (UTRs). HCV replication requires the presence of a host microRNA (miRNA), miR-122, the most abundant miRNA in hepatocytes (Jopling, Yi, Lancaster, Lemon, & Sarnow, 2005). The miR-122 dependency of HCV partly explains its hepatic tropism.

In infected hepatocytes, miR-122 binds to two specific sites located within the HCV 5’-UTR, thereby playing a major role in the viral lifecycle. The 5’-UTR contains the internal ribosome entry site (IRES) that mediates cap-independent translation of viral proteins (Friebe, Lohmann, Krieger, & Bartenschlager, 2001; Honda, Beard, Ping, & Lemon, 1999). miR-122 binding to the 5’-UTR triggers AGO2 recruitment, thereby protecting the HCV RNA molecules from 5’-RNA decay (Amador-Canizares, Bernier, Wilson, & Sagan, 2018; Li, Yamane, Masaki, & Lemon, 2015; Sedano & Sarnow, 2014; Shimakami et al., 2012). Finally, the Poly(RC)-binding protein 2 (PCBP2) is displaced from the viral RNA by miR-122, resulting in increased replication and decreased translation (Masaki et al., 2015). Whether and how HCV itself regulates the maturation of miR-122, thereby controlling its own lifecycle, remains unexplored.

KH-type splicing regulatory protein (KSRP) is a ubiquitous nuclear and cytoplasmic RNA-binding protein that plays a key role in miRNA maturation. Indeed, KSRP is a component of DROSHA and Dicer complexes, which contribute to the maturation of a class of miRNAs in both the nucleus and cytosol (Repetto et al., 2012; Trabucchi et al., 2009). KSRP is also involved in the degradation of AU-rich element (ARE)-containing cellular mRNAs (Briata, Chen, Ramos, & Gherzi, 2013; Gherzi, Chen, Trabucchi, Ramos, & Briata, 2010; Kroll, Zhao, Jiang, & Huber, 2002; Min, Turck, Nikolic, & Black, 1997). We previously observed that ARE-containing mRNAs are upregulated in HCV-infected hepatocytes, a result suggesting that this function of KSRP could be altered during viral infection (Colman et al., 2013). KSRP is phosphorylated at 5 residues: by mitogen-activated protein kinase (MAPK) p38 at threonine residue 692; by ataxia-telangectasia-mutated (ATM) at serine residues 132, 274 and 670; and/or by Akt at serine residue 193. The Akt-dependent phosphorylation of KSRP at serine 193 promotes nuclear pri-miRNA processing in the context of reduced ARE-containing mRNA degradation (Briata et al., 2013; Briata et al., 2005; Y. Liu & Liu, 2011; Zhang, Wan, Berger, He, & Lu, 2011).

HCV, particularly its non-structural 5A (NS5A) protein, has been shown to activate the phosphatidylinositol-3-kinase (PI3K) signaling pathway in many models, thereby activating its downstream effector, the serine/threonine kinase Akt (Bose, Meyer, Di Bisceglie, Ray, & Ray, 2012; Cheng et al., 2015; Han, Niu, Wang, & Li, 2016; He et al., 2002; Higgs, Lerat, & Pawlotsky, 2013; Milward, Mankouri, & Harris, 2010; Street, Macdonald, Crowder, & Harris, 2004). Here, we demonstrate that HCV infection induces KSRP phosphorylation at serine position 193 through Akt activation and that this P-S193-KSRP in turn induces the maturation of miR-122, which favors viral replication.

## Results

### KSRP exerts a proviral effect on HCV replication mediated by miR-122 maturation

Because the ubiquitous cellular protein KSRP is involved in the processing of miRNAs in HeLa cells (Trabucchi et al., 2009), we assessed whether maturation of miR-122, the most abundant liver-specific miRNA and a key actor of the HCV lifecycle, is KSRP-dependent. miR-23b, another less abundant miRNA present in hepatocytes but not involved in the HCV lifecycle, was used as a control. KSRP was silenced by means of siRNA transfection in non-infected Huh7.5 cells and total RNA and proteins were analyzed at different time points. Transient KSRP silencing led to a significant decrease of the intracellular amount of both miR-122 and miR-23b at 48 and 72 hours post-transfection (Figure 1A). We confirmed by western blot analysis the decrease of KSRP protein quantity as compared to the negative silencing control and the lack of cytotoxicity of KSRP silencing at the selected time points (Supplementary Figure S1).

**Figure 1.**
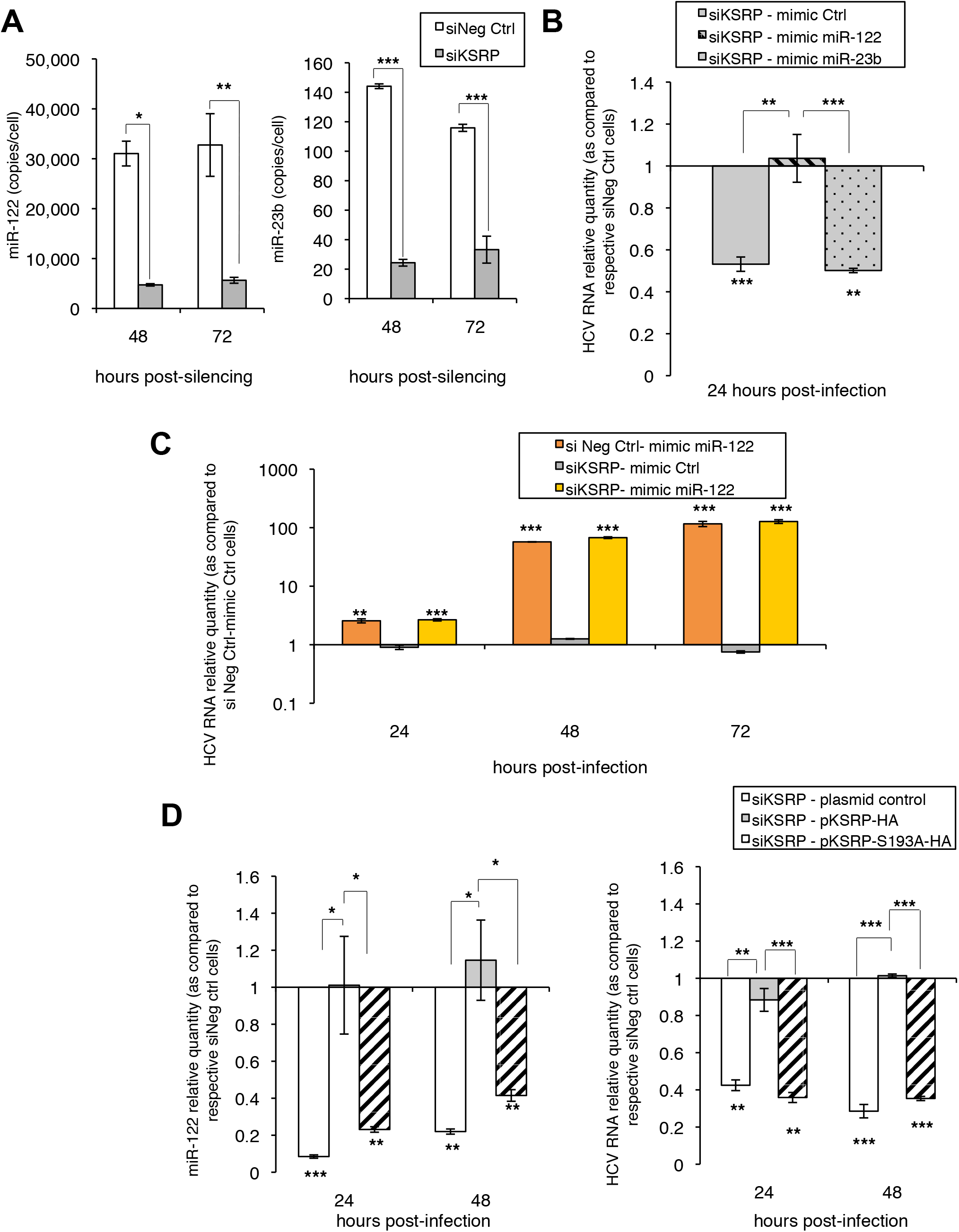
Effect of KSRP on miR-122 maturation and HCV replication. **(A)** RT-qPCR quantification of miR-122 and miR-23b (control miRNA with no role in the HCV lifecycle) in KSRP-silenced (siKSRP) and non-silenced (siNeg Ctrl) Huh7.5 cells. The miRNA copy numbers per cell at 48 and 72 h post-siRNA transfection are shown as mean±SEM from three replicate experiments. **(B)** Rescue of HCV replication in KSRP-silenced cells (siKSRP) by mimic miR-122 and mimic miR-23b transfection, as assessed by RT-qPCR in comparison with a negative control mimic miRNA (mimic Ctrl). HCV RNA quantities were normalized to GAPDH mRNA and expressed as mean±SEM from three replicate experiments to the HCV RNA amount in non-silenced cells transfected with the negative mimic miRNA control (siNeg Ctrl-mimic Ctrl). **(C)** Kinetics of HCV RNA levels, as assessed by RT-qPCR, in Hep3B cells co-transfected with siKSRP or a negative silencing control (siNeg Ctrl) and a mimic miR-122 or a negative mimic miRNA control (mimic Ctrl), and infected 24 h post-transfection with the HCV Jad strain at an MOI of 0.1. HCV RNA quantities were normalized to GAPDH mRNA and expressed as mean±SEM from three replicate experiments to the HCV RNA amount in non-silenced cells transfected with the negative mimic miRNA control (siNeg Ctrl-mimic Ctrl). **(D)** miR-122 and HCV RNA levels assessed by RT-qPCR quantification in Huh7.5 cells silenced for KSRP by means of siKSRP^e^ treatment, subsequently transfected with plasmids expressing KSRP-HA (pKSRP-HA) or non-phosphorylable KSRP-S193A-HA (pKSRP-S193A-HA) or a negative plasmid control, then infected with the HCV Jad strain at an MOI of 0.1. miR-122 and HCV RNA quantities were normalized to RNU6B (U6) or GAPDH mRNA, respectively, and are expressed as mean±SEM from three replicate experiments to the quantities in cells treated with the negative silencing control transfected with the negative plasmid control (siNeg Ctrl – plasmid control).

The role of KSRP on HCV infection was then studied. For this, Huh7.5 cells treated with siKSRP or the negative silencing control were cotransfected with exogenous mimic miR-122, mimic miR-23b (control miRNA) or a negative mimic control. Twenty-four hours post-transfection, the cells were infected with the HCV Jad strain at an MOI of 0.1 and HCV RNA was quantified 24 h post-infection. As shown in Figure 1B, HCV replication was significantly reduced 24 h post-infection in siKSRP-treated Huh7.5 cells as compared to the negative control. The exogenous addition of mimic miR-122 restored HCV replication in KSRP-deficient Huh7.5 cells, whereas mimic miR-23b or the negative mimic control did not (Figure 1B). These results parallel the effects of transient KSRP silencing and mimic miRNA co-transfections on miR-122 and miR-23b levels, as assessed by RT-qPCR and on KSRP amounts as assessed by western blot analysis, in HCV-infected Huh7.5 cells (Supplementary Figure S2A and S2B).

To confirm these results, we used Hep3B cells, hepatoma-derived cells with undetectable levels of miR-122, in which HCV replication and the production of viral particles are fully dependent on mimic miR-122 supplementation (Thibault, Huys, Dhillon, & Wilson, 2013; Varnholt et al., 2008). Hep3B cells were co-transfected with siKSRP or the negative silencing control on the one hand, mimic miR-122 or the mimic control on the other hand, 24 h before HCV infection with the Jad strain at an MOI of 0.1 (the experimental design is shown in the Supplementary Figure S3A). The respective amounts of miR-122 and KSRP protein obtained after co-transfection in Hep3B cells are shown in the Supplementary Figure S3B and S3C. As shown in Figure 1C, HCV replication was restored in Hep3B cells by mimic miR-122 transfection and it was not altered when KSRP was silenced in miR-122-co-transfected cells.

Together, these results suggest that KSRP plays a proviral role in HCV-infected hepatocytes by promoting the maturation of miR-122, a key actor of the HCV lifecycle.

### The proviral role of KSRP in HCV infection requires its phosphorylation at serine residue 193

Akt-dependent phosphorylation of KSRP at serine residue 193 has been reported to promote miRNA maturation by facilitating pri-miRNA to pre-miRNA processing (Briata et al., 2013; Briata et al., 2012). Therefore, we investigated the effect of non-phosphorylable KSRP-S193A mutagenesis on miR-122 quantities and HCV replication. Huh7.5 cells were transfected with a pool of 3 siRNAs targeting the 3’- UTR region of KSRP mRNA (siKSRP^e^) to block the production of endogenous KSRP. Then, plasmids expressing exogenous HA-tagged wild-type KSRP or KSRP-S193A were transfected for 48 h before HCV infection of Huh7.5 cells at an MOI of 0.1. HCV replication and miR-122 amounts were quantified by means of RT-qPCR at 24 h and 48 h post-infection (see experimental design and the quantities of KSRP and KSRP-HA proteins obtained 48 h post-infection in the Supplementary Figure S4). As shown in Figure 1D, the reductions of miR-122 and HCV RNA quantities induced by KSRP silencing at 24 and 48 h were reversed by transfection of pKSRP-HA but not with non-phosphorylable pKSRP-S193A-HA or the plasmid control. These results suggest that the phosphorylation of KSRP at serine residue 193 is essential for miR-122 maturation and HCV replication.

### KSRP phosphorylation at serine residue 193 is increased in the presence of HCV replication

We then aimed to study the Akt-dependent phosphorylation of S193-KSRP in Huh7.5 cells 24 h post-infection by the genotype 2a JFH1-derived HCV strain Jad at an MOI of 0.1. Because no specific antibodies against the phosphorylated forms of KSRP are currently available, we quantified and analyzed the subcellular localization of S193-phosphorylated KSRP by means of *in situ* proximity ligation assay (PLA) using anti-KSRP^mouse^ and anti-RXXS*/T* primary antibodies. As shown in Figure 2A, *in situ* PLA showed markedly increased amounts of both cytoplasmic and nuclear phospho-S193-KSRP in Jad-infected Huh7.5 cells as compared to uninfected cells. Control experiment without primary antibodies is shown in the Supplementary Figure S5.

**Figure 2.**
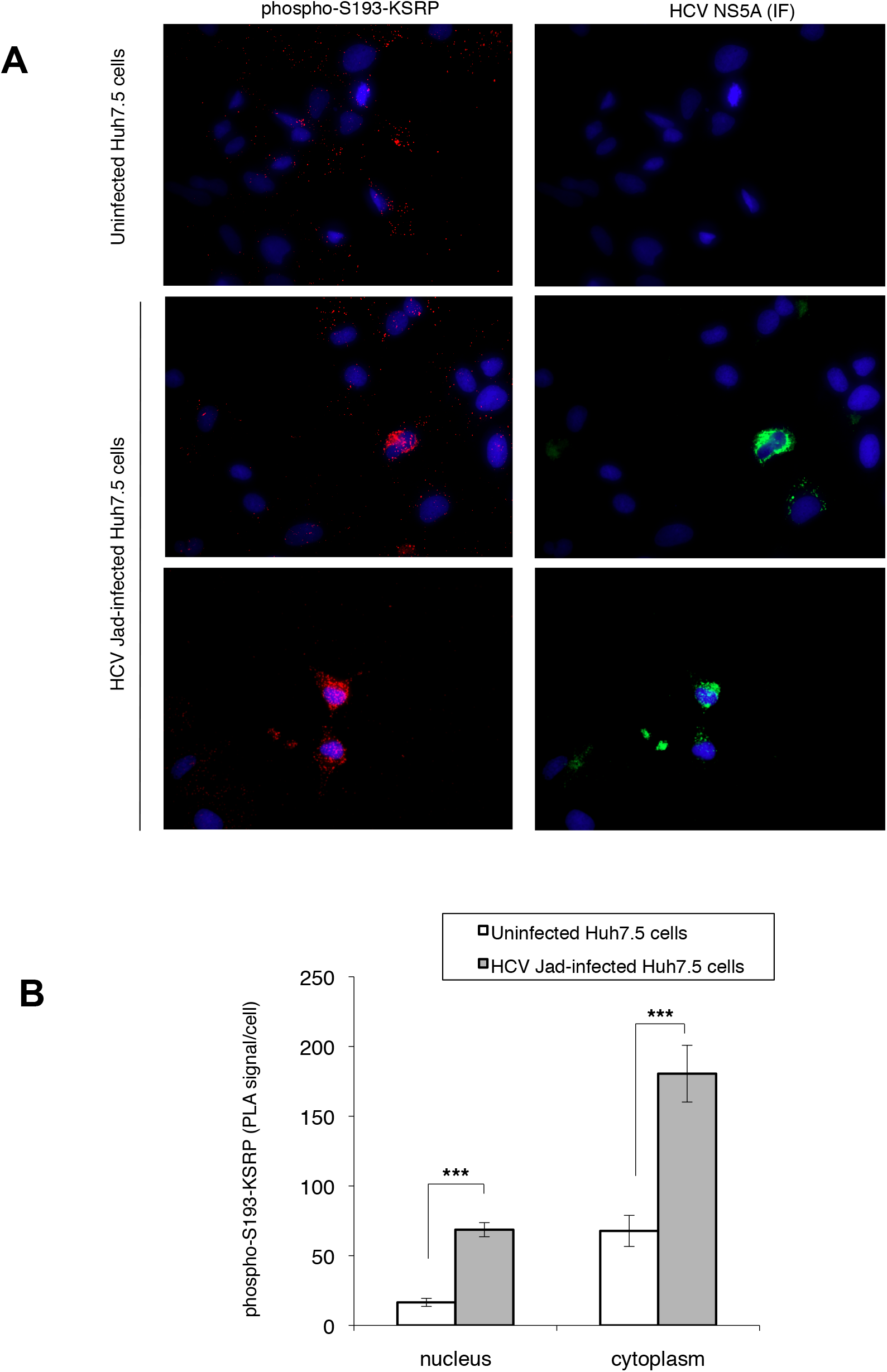
Effect of HCV infection on the amount of nuclear and cytoplasmic phospho-S193-KSRP in hepatoma cell line, as assessed by *in situ* PLA. **(A)** Representative *in situ* PLA images showing nuclear and cytoplasmic phospho-S193-KSRP in Huh7.5 cells infected with the HCV genotype 2a Jad strain at an MOI of 0.1, as compared to uninfected cells. **(B)** Signal quantification of *in situ* PLA experiments obtained using the Duolink Image Tool software. In **(A)**, cells were probed with antibodies directed against the HCV NS5A protein (immunofluorescence, IF signal, green) or phospho-S193-KSRP (KSRP^mouse^ x RXXS*/T*) (PLA signal, red). Cell nuclei were stained with DAPI (blue). In **(B**), the amounts of phospho-S193-KSRP are expressed as mean±SEM of PLA signals per cell. Values are calculated from n=7-30 cells.

### Phosphorylation of Akt at serine 473 is induced by HCV infection, is required for the phosphorylation of KSRP at serine 193 and for HCV replication

To assess whether the increased quantity of phospho-S193-KSRP observed in presence of HCV could result from a viral-induced activation of the kinase Akt, we first analyzed by western blot the phosphorylation of Akt at position 473, 32 hours post infection of Huh7.5 cells by the Jad strain of HCV. As shown in Figures 3A and 3B, HCV infection was associated with the activation of the PI3K/Akt pathway through the phosphorylation of Akt at position 473.

**Figure 3.**
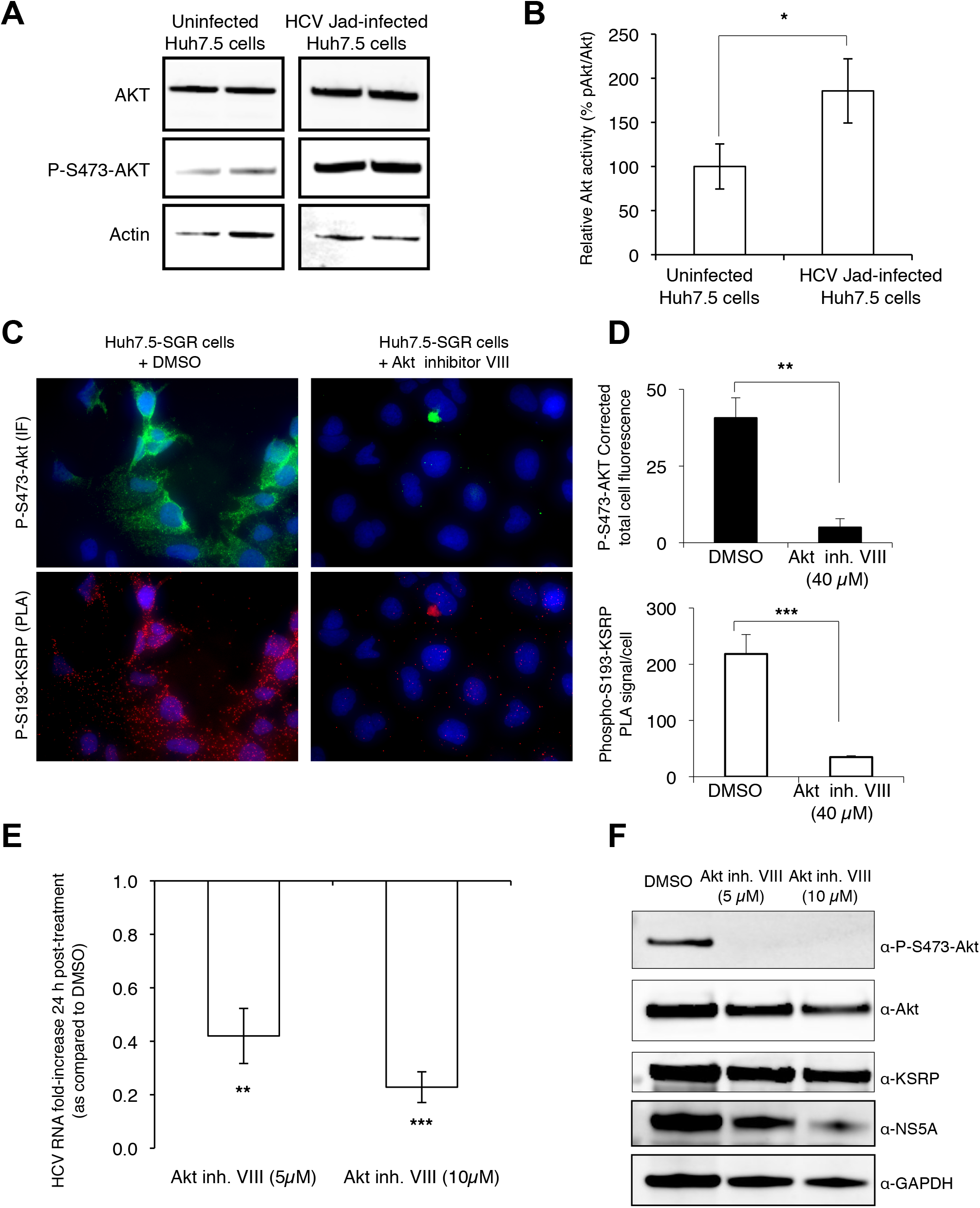
HCV infection favors S473-Akt phosphorylation which is required for S193-KSRP phosphorylation and HCV replication. **(A)** Western blot analysis of Akt and phospho-S473-Akt in uninfected Huh7.5 cells and in cells infected with the HCV genotype 2a Jad strain at an MOI of 0.1. Huh7.5 cells were starved in DMEM-low glucose (1g/l) medium for 16 h, then infected with Jad for 8 h, washed with PBS and cultured in DMEM-low glucose medium for 24 h. **(B)** Histogram representing the percentage expressed as mean±SEM from two replicate experiments of phospho-S473-Akt relative to total Akt in uninfected Huh7.5 cells and in cells infected with the HCV genotype 2a Jad strain at an MOI of 0.1. Quantifications were performed using the luminescence image analyzer Image Quant Las 4000 mini and Image Quant software (GE Healthcare). **(C)** Representative images showing phospho-S193-KSRP (*in situ* PLA) and phospho-S473-Akt (immunofluorescence, IF) in the absence (DMSO) or in the presence of 40 !M Akt inhibitor VIII administered for 6 h in Huh7.5 cells harboring the HCV genotype 1b subgenomic replicon (Huh7.5-SGR). Cells were probed with antibodies directed against phospho-S473-Akt antibody (IF signal, green) and phospho-S193-KSRP (KSRP^mouse^ x RXXS*/T*) (PLA signal, red). Cell nuclei were stained with DAPI (blue). **(D)** Top: quantification of phospho-S473-Akt levels (IF signals) in Huh7.5-SGR cells in the absence (DMSO) or in the presence of 40 !M Akt inhibitor VIII by means of ImageJ software (v1.48, NIH). The results are expressed as mean±SEM of corrected total cellular fluorescence (CTCF=Integrated Density – (Area of selected cell X Mean fluorescence of background readings) per cell (n=7 cells). Bottom: quantification of phospho-S193-KSRP levels (PLA signals) in Huh7.5-SGR cells in the absence (DMSO) or in the presence of 40 !M Akt inhibitor VIII by means of the Duolink Image Tool software. The results are expressed as mean±SEM of total PLA signals per cell (n=7 cells). **(E)** Quantification of HCV replicon RNA levels by RT-qPCR in Huh7.5 in the presence of 5 !M or 10 !M of Akt inhibitor VIII administered for 24 h using GAPDH as endogenous control relative to DMSO. Results are expressed as mean±SEM from three replicate experiments. **(F)** Western blot analysis showing the effect of 5 !M or 10 !M Akt inhibitor VIII administered for 24 h on Akt, phospho-S473-Akt, KSRP and HCV NS5A protein expression in Huh7.5-SGR cells. GAPDH was used as the loading control.

Then, we tested the effect of Akt inhibition by Akt inhibitor VIII on S193-KSRP phosphorylation by means of *in situ* PLA experiments using anti-KSRP^mouse^ and anti-RXXS*/T* primary antibodies coupled with an immunofluorescence assay using specific antibodies to detect phospho-S473-Akt, in Huh7.5 cells harboring the HCV genotype 1b subgenomic replicon. As shown in Figures 3C and 3D, the amounts of both phospho-S193-KSRP and phospho-S473-Akt were significantly reduced by Akt inhibitor VIII treatment in HCV replicon-harboring cells, as compared to untreated cells. These results confirm the dependency of S193-KSRP phosphorylation for Akt activity.

We further analyzed the effects of the inhibition of Akt phosphorylation at serine 473 by Akt inhibitor VIII on HCV replication and viral protein expression. Inhibition of Akt activity was associated with a decrease of the amount of HCV replicon RNA and NS5A protein in replicon-harboring cells, whereas the total amount of KSRP protein remained unchanged (Figures 3E and 3F).

These findings demonstrate a viral-induced activation of Akt, as measured by the amount of phosphor-S473-Akt and show a correlation between Akt activity and S193-KSRP phosphorylation. Finally, our results emphasize the requirement of Akt activity for HCV replication and protein expression.

### Akt-dependent phosphorylation of S193-KSRP induces miR-122 maturation and enhances HCV replication

As shown in Figure 4A, Akt inhibitor VIII treatment of Huh7.5 cells harboring the HCV genotype 1b subgenomic replicon was also associated with significant dose-dependent increase of the amount of pri-miR-122 and decrease of the amount of mature miR-122. This result is in keeping with the requirement of phospho-S193-KSRP for miR-122 maturation. In light of these findings, we hypothesized that the diminution of HCV replication in Akt inhibitor VIII treated-cells (Figure 3E and 3F) was linked to the reduced phosphorylation of S193-KSRP (Figure 3C and 3D) and thus, to the decrease of miR-122 maturation (Figure 4A).

**Figure 4.**
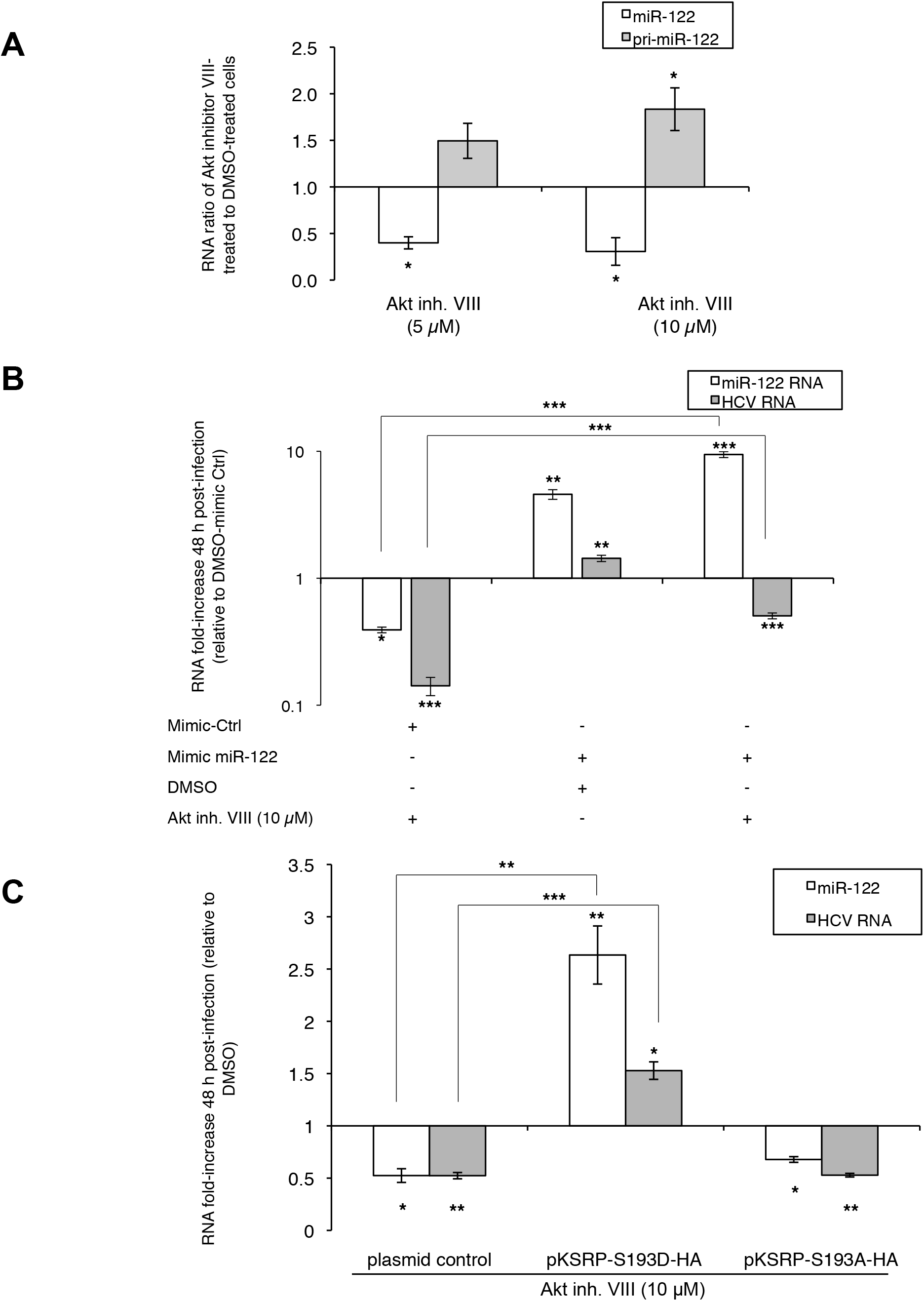
Role of Akt-dependent phosphorylation of KSRP at serine 193 in miR-122 maturation and HCV replication. **(A)** Effect of Akt inhibition on pri-mir-122 and miR-122 quantities, as measured by RT-qPCR in Huh7.5 cells harboring the HCV genotype 1b subgenomic replicon. The results are shown as the RNA quantity ratio in Akt inhibitor-treated cells relative to DMSO-treated cells. Pri-miR-122 and miR-122 quantities were normalized to GAPDH and RNU6B (U6) RNAs, respectively. Error bars represent SEM from three replicate experiments. **(B)** Mature miR-122 and HCV RNA quantities in Huh7.5 cells treated with 10 !M of Akt inhibitor VIII or DMSO and transfected with a negative mimic control (mimic Ctrl) or a mimic miR-122, as assessed by RT-qPCR. miR-122 and HCV RNA quantities were normalized to RNU6B (U6) and GAPDH, respectively, and are presented as mean±SEM from three replicate experiments to the corresponding quantities in cells treated with DMSO and mimic Ctrl. **(C)** Mature miR-122 and HCV RNA quantities in Huh7.5 cells treated with 10 !M of Akt inhibitor VIII or DMSO and transfected with plasmids expressing different forms of KSRP tagged with HA, as assessed by RT-qPCR. miR-122 and HCV RNA quantities were normalized to RNU6B (U6) and GAPDH, respectively, and are presented as mean±SEM from three replicate experiments to the corresponding quantities in cells treated with DMSO and transfected with the control plasmid.

To evaluate this hypothesis, Huh7.5 cells were pre-incubated with DMSO or 10 !M Akt inhibitor VIII for 8 h, transfected with mimic Ctrl or mimic miR-122 and infected 24 h post-treatment with the HCV Jad strain at an MOI of 0.1. Akt inhibitor VIII treatment or DMSO was maintained for the total duration of the experiment, i.e. until 48 h post-infection. As shown in Figure 4B, there was a 2.5-fold decrease of miR-122 and a 8.3-fold decrease of HCV RNA quantities in cells treated with Akt inhibitor VIII in the absence of mimic-miR-122, as compared to the DMSO-treated control. The amount of miR-122 was increased by the administration of exogenous mimic miR-122 in both DMSO-treated and Akt inhibitor VIII-treated cells. An exogenous mimic miR-122 transfection to Akt inhibitor VIII-treated cells restored a level of HCV replication close to that observed in DMSO- and mimic control-treated cells, but slightly lower than that in DMSO- and mimic miR-122-treated cells (Figure 4B). These results suggest that the effect of Akt inhibition on HCV replication is at least partly related to a reduction of the amount of hepatocyte miR-122 as a result of a defect in its maturation.

We then assessed whether the antiviral effect of Akt inhibitor VIII could be reversed by KSRP transfection. Huh7.5 cells were pre-treated for 8 h with 10 μM of Akt inhibitor VIII or DMSO, transfected with KSRP-HA plasmids for 16 h and infected with the Jad strain of HCV. Again, Akt inhibitor VIII or DMSO treatment was maintained for the total duration of the experiment, i.e. until 48 h post-infection. As shown in Figure 4C, Akt inhibitor VIII reduced the amounts of miR-122 and HCV RNA in control cells and in cells transfected with non-phosphorylable KSRP-S193A-HA. In contrast, transfection of a plasmid expressing phosphomimetic KSRP-S193D-HA rescued both miR-122 quantities and HCV replication in the presence of Akt inhibitor VIII (Figure 4C).

These results suggest that Akt-dependent phosphorylation of KSRP at serine 193 induces the maturation of miR-122, thereby favoring HCV replication.

### Phospho-S193-KSRP interacts with pri-miR-122 and DROSHA

Because Akt-dependent phosphorylation of S193-KSRP is involved in the processing of pri-miRNA to pre-miRNA (Briata et al., 2013; Briata et al., 2012), we investigated the potential interaction between KSRP and pri-miR-122. RNA immunoprecipitation experiments using antibodies against KSRP and DROSHA showed co-immunoprecipitation of both proteins with pri-miR-122. Immunoprecipitation of pri-miR-122 with an anti-DROSHA antibody was significantly reduced by transient silencing of KSRP in Huh7.5 cells harboring the HCV genotype 1b subgenomic replicon (Figure 5A). Similar experiments were conducted with HA-tagged wild-type and S193-mutated KSRP using an anti-HA antibody. Pri-miR-122 interacted with both KSRP-HA and phosphomimetic KSRP-S193D-HA (11.3- and 81.8-fold increase in pri-miR-122 amount in the immunoprecipitate, respectively), but not with non-phosphorylable KSRP-S193A-HA. Control experiments performed with c-jun ARE mRNA showed a strong interaction with KSRP-HA and KSRP-S193A-HA proteins (33.0- and 40.6-fold increase in c-jun mRNA amount in the immunoprecipitate, respectively) (Figure 5B), indicating that non-phosphorylated S193-KSRP interacts with ARE-containing cellular mRNAs.

**Figure 5.**
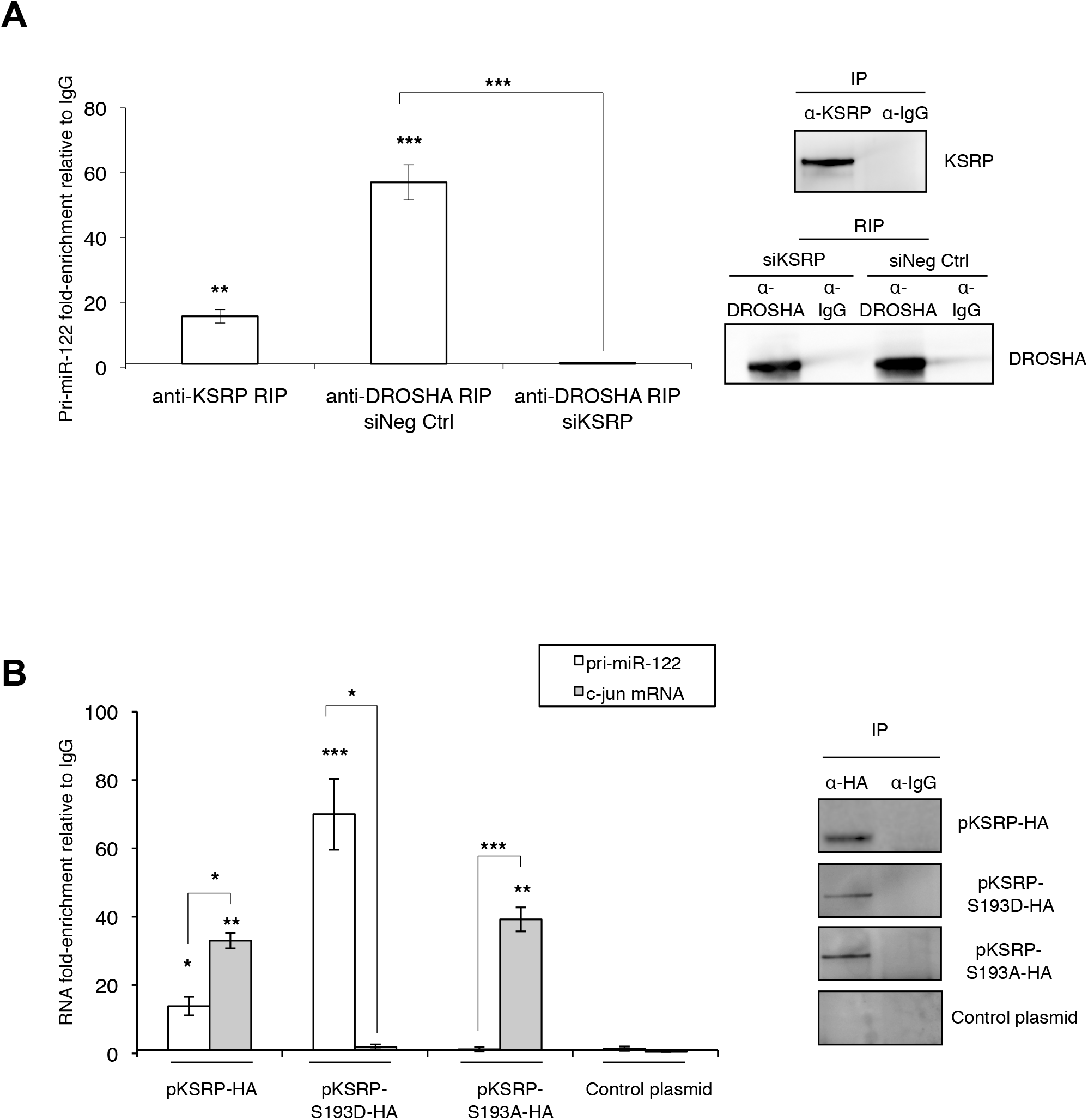
Role of KSRP in the formation of the DROSHA/pri-miR-122 complex. **(A)** RNA immunoprecipitation (RIP) experiments showing the interaction between KSRP, pri-miR-122 and DROSHA and the effect of KSRP silencing on this interaction in the nucleus of Huh7.5 cells harboring the HCV subgenomic replicon. Pri-miR-122 enrichment in the immunoprecipitate was measured by RT-qPCR, normalized to input RNA and quantified relative to an isotype control IgG antibody (left). Results are expressed as mean±SEM from three replicate experiments. Immunoblot analysis of the immunoprecipation fractions using anti-KSRP, anti-DROSHA and anti-IgG antibodies (right). **(B)** RNA immunoprecipitation (RIP) experiments showing the interaction between wild-type and mutated HA-tagged KSRP proteins, with nuclear pri-miR-122 and cytoplasmic c-jun mRNA in Huh7.5 cells harboring the HCV subgenomic replicon. Pri-miR-122 and c-jun mRNA enrichment in the immunoprecipitate was measured by RT-qPCR, normalized to input RNA and quantified relative to an isotype control IgG antibody (left). Results are expressed as mean±SEM from three replicate experiments. Immunoblot analysis of the immunoprecipation fractions using anti-HA and anti-IgG antibodies (right).

These findings suggest that Akt-dependent phosphorylation of KSRP at serine 193 is required for the interaction of pri-miR-122 with DROSHA that leads to pri-miR-122 maturation.

### Akt activation and KSRP expression are required to maturate miR-122 and promote HCV replication in primary human hepatocytes

Because primary human hepatocytes (PHHs) are the most relevant cellular model for the study of HCV infection, previous experiments were replicated in PHHs infected with a non-modified genotype 2a JFH1 virus at a MOI of 2. Treatment with 4 !M of Akt inhibitor VIII for 96 h reduced Akt phosphorylation by 55.3%, as shown by western blot analysis (Figure 6A). Akt inhibitor VIII treatment was also associated with an approximately 2-fold decrease of HCV RNA and mature miR-122 amounts and no change in KSRP mRNA expression (Figure 6B). KSRP silencing, performed 24 h before JFH1 infection of PHHs at a MOI of 2, was associated with a marked reduction of both miR-122 expression and HCV replication as compared to non-silenced PHHs (Figures 6C and 6D).

**Figure 6.**
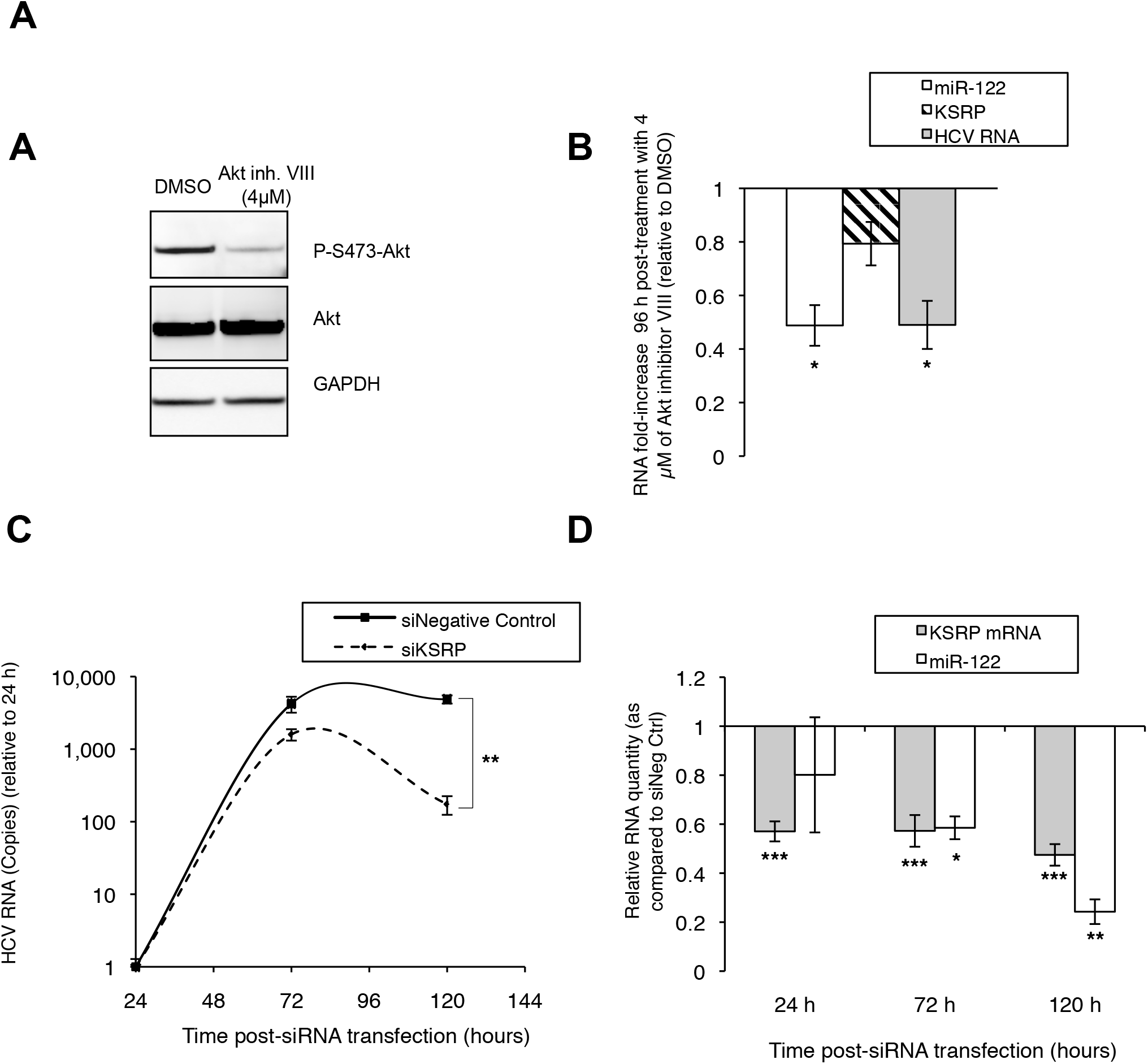
Effect of Akt inhibition and KSRP silencing in primary human hepatocytes infected with the unmodified JFH1 HCV strain. **(A)** Western blot analysis showing the effect of Akt inhibitor VIII on phospho-S473-Akt expression. GAPDH was used as the loading control. **(B)** Effect of Akt inhibitor VIII on HCV replication (as assessed by quantification of intracellular HCV RNA amounts) and on the expression of miR-122 and KSRP mRNA (as assessed by RT-qPCR). RNA quantities were normalized to GAPDH or RNU6B (U6) and expressed as mean±SEM from three replicate experiments. **(C)** Kinetics of HCV replication, as assessed by quantification of intracellular HCV RNA by RT-qPCR in PHHs silenced for KSRP expression, as compared to non-silenced PHHs (siNegative Control) starting 24 h post siRNA transfection. HCV RNA quantity was normalized to GAPDH and expressed as mean±SEM from three replicate experiments. **(D)** Kinetics of KSRP and miR-122 RNA expression in PHHs silenced for KSRP expression, as compared to non-silenced PHHs (siNegative Control). RNA quantities were normalized with GAPDH or RNU6B (U6) and expressed as mean±SEM from three replicate experiments.

These results confirm the essential role of Akt activity and KSRP in miR-122 maturation and HCV replication in HCV-infected PHHs.

## Discussion

miR-122 is the most abundant tissue-specific miRNA in hepatocytes and is required to achieve a full HCV lifecycle. This requirement partly explains the hepatic tropism of HCV. Indeed, the ectopic expression of miR-122 and apolipoprotein E (ApoE) has been shown to be sufficient to render a mouse liver-derived cell line permissive for HCV replication and for the release of infectious particles (Frentzen et al., 2014). Additionally, the expression of exogenous miR-122 in non-hepatic or hepatic cell lines facilitates efficient HCV replication. The susceptibility of Huh7.5 cells to the propagation of infectious HCV strains in cell culture is attributable to their high level of miR-122 expression (Fukuhara et al., 2012).

Binding of miR-122 at two tandem sites, S1 and S2, located within the 5’ untranslated region of the viral genome enhances RNA stability and promotes viral replication (Jopling, Schutz, & Sarnow, 2008; Li, Masaki, & Lemon, 2013). Until now, whether and how HCV can regulate the maturation of miR-122 to sustain its own lifecycle remained unexplored. Our results show that the host protein KSRP is essential for miR-122 maturation in hepatocytes (Figure 1A) and therefore is a proviral host factor for HCV (Figures 1B). We also demonstrate that Akt phosphorylation of S193-KSRP is pivotal for this function (Figures 1D). In the present work, we used four cell culture-based models, including hepatoma-derived cells harboring a stable genotype 1b subgenomic replicon, Hep3B and Huh7.5 cells infected with the genotype 2a JFH1-derived strain Jad, and PHHs infected with the unmodified genotype 2a JFH1 strain, to uncover the existence of a reciprocal regulation loop where HCV infection could modulate miR-122 maturation, and therefore fuel its own replication. This pathway consists of several sequential steps : (i) HCV infection activates Akt by phosphorylation at serine residue 473 (Figure 3A); (ii) activated Akt in turn phosphorylates KSRP at serine residue 193 (Figures 3C-D), resulting in increased amounts of phospho-S193-KSRP (Figure 2); (iii) phospho-S193-KSRP binds to nuclear pri-miR-122 (Figure 5B), and KSRP promotes the association between pri-miR-122 with DROSHA (Figure 5A); (iv) phospho-S193-KSRP and DROSHA binding induces pri-miR-122 maturation into pre-miR-122 within this DROSHA complex, then maturation of pre-miR-122 into miR-122; (v) Akt dependent phosphorylation of S193-KSRP favors miR-122 expression in hepatocytes which in turn stimulates HCV replication (Figure 4C). Interestingly, our results indicate a complete rescue of HCV replication in Akt VIII inhibitor treated-cells after transfection with a plasmid expressing phosphomimetic S193D-KSRP but not after mimic-miR-122 transfection (Figures 4B-C). Indeed, in the context of Akt inhibition, transfection of significant amount of mimic-miR-122 does not seem to fully restore HCV replication (Figure 4B, lane mimic-miR-122 and Akt inh. VIII). These results suggest an additional proviral function for S193D-KSRP, possibly linked to the maturation of other miRNAs than miR-122 favoring, directly or indirectly, the HCV lifecycle. This original, thus far unknown model of self-regulation of the HCV lifecycle involving the cellular protein KSRP is summarized in Figure 7.

**Figure 7.**
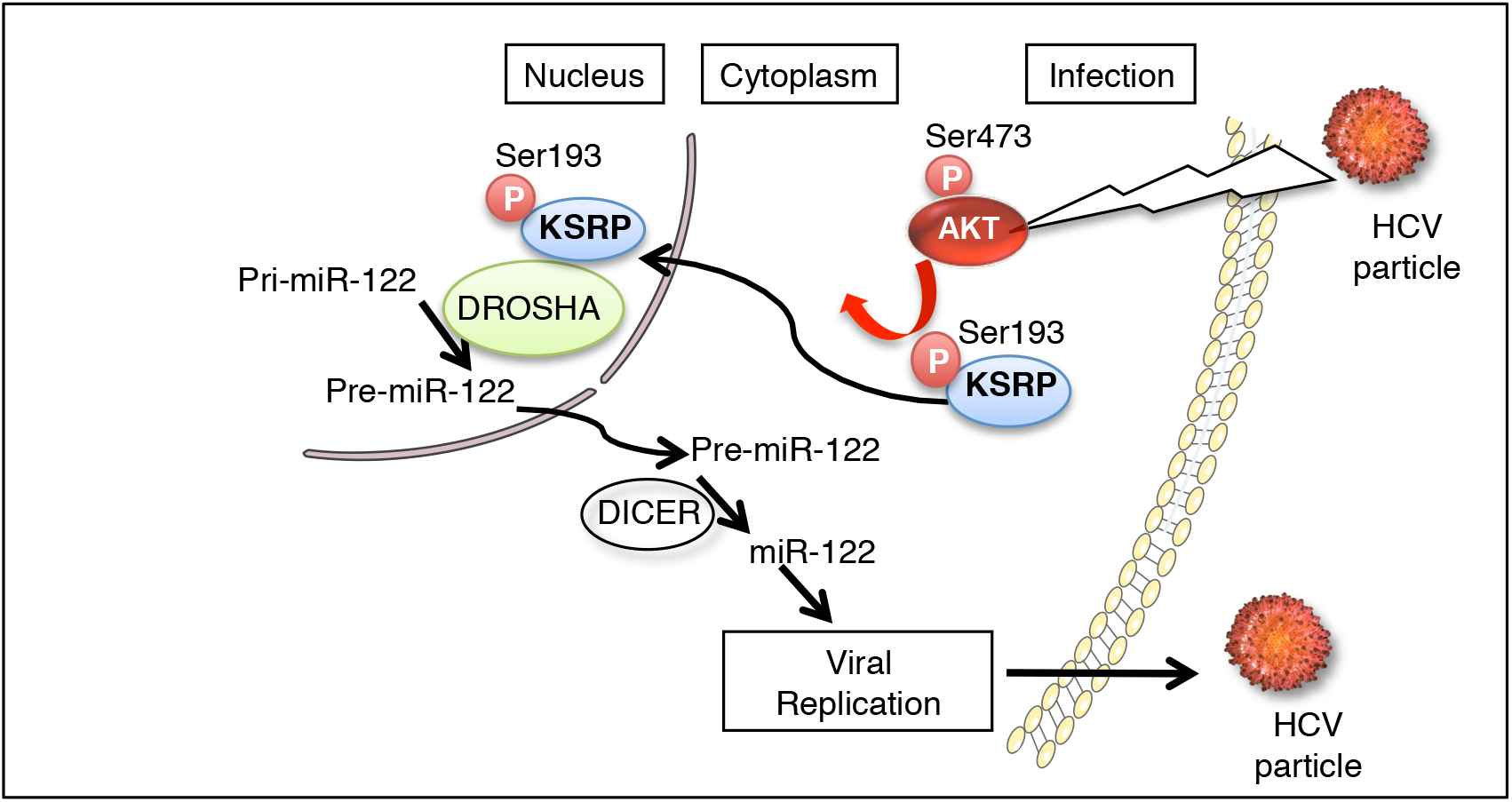
Proposed model of HCV-induced, Akt-dependent activation of KSRP resulting in miR-122-mediated induction of viral replication. HCV infection activates Akt through its phosphorylation at serine residue 473; phospho-S473-Akt phosphorylates KSRP at serine residue 193, resulting in increased amounts of phospho-S193-KSRP in the nucleus where phospho-S193-KSRP binds pri-miR-122 and DROSHA; phospho-S193-KSRP and DROSHA binding induces pri-miR-122 maturation into pre-miR-122 and the release of pre-miR-122 in the cytoplasm where it is in turn matured into miR-122; miR-122 stabilizes HCV RNA and induces HCV replication and translation through already described mechanisms, ultimately leading to the release of infectious viral particles after subsequent steps of the HCV lifecycle have been completed.

Our results showing that HCV-induced maturation of miR-122 is mediated by KSRP as a result of Akt activation *via* its phosphorylation at serine residue 473 is in keeping with previous observations of the activation of PI3K/Akt signaling pathway during HCV infection (Z. Liu et al., 2012). Accumulating evidence suggests that the NS5A protein, alone or in association with other non-structural viral proteins, plays an important role in HCV-induced Akt activation (Cheng et al., 2015; Han et al., 2016; Higgs et al., 2013; Shi, Hoffman, & Liu, 2016). Altogether, our results show that, in addition to its many reported effects in HCV-infected cells, HCV-induced Akt activation plays a key role in the viral lifecycle by regulating replication through miR-122 maturation. Moreover, our observation of an increased phosphorylation of S193-KSRP *via* the activated PI3K/Akt signaling pathway resulting in miRNA maturation and reduced ARE mRNAs interaction in HCV-infected cells (Figure 5B) is also in keeping with a recent report of an upregulation of ARE-dependent mRNAs at day 3 post-infection with JFH1 (Colman et al., 2013).

Previous studies indicated that miR-122 levels are decreased in cells and in sera from patients chronically infected with HCV (Bala, Marcos, & Szabo, 2009; Marquez et al., 2010). These observations could be linked to the decreased stability of miR-122 resulting from the interaction between HCV core protein and GLD-2, leading to the inhibition of this non-canonical cytoplasmic poly(A) polymerase (Kim et al., 2016). In addition, miR-122 “sponging” by HCV RNA has been reported during HCV infection. Indeed, HCV RNA acts as a competitive inhibitor of miR-122 activity by establishing the stable interactions required for viral genome stability and replication. Because HCV RNA is a template for its replication, a positive feedback loop sequesters additional miR-122. This “sponging” phenomenon has the potential to result in a global de-repression of cellular miR-122 target genes (Luna et al., 2015). Therefore, compensatory mechanisms are required to maintain the cellular levels of miR-122 while sustaining both HCV RNA replication and post-translational regulation of miR-122 cellular target genes. Thus, the regulation of miR-122 maturation by KSRP described here could be of importance for acute HCV infection and the long-term homeostasis of infected hepatocytes in the context of chronic HCV infection.

In summary, we describe here a so far unknown mechanism by which HCV “hijacks” KSRP, a host cell protein implicated in RNA metabolism, to regulate the hepatocyte expression of miR-122 and therefore fuel its own replication. Whether this “hijacking” is specific for HCV infection or could represent a general mechanism used by different viruses to establish and sustain their chronic infection remains to be investigated. miR-122 has been shown to be an attractive target for HCV inhibition. Indeed, a single injection of an antagomir drastically reduced viral replication for several weeks, leading to a cure of infection in some cases (van der Ree et al., 2017). These results validated the concept that antagonists or agonists of miRNAs with proviral or antiviral effects, respectively, could be used in the treatment of acute or chronic viral infections. Our results suggest that KSRP may constitute a valuable tool to identify new targets of antiviral intervention.

## Materials and Methods

### Cell lines

Three hepatoma cell lines were used: Hep3B cells; Huh7.5-SGR cells, i.e. Huh7.5 cells stably harboring the neomycin resistance gene and a genotype 1b HCV subgenomic replicon derived from plasmid I389-Neo/NS3-3!/5.1 (kindly provided by Dr Ralf Bartenschlager, University of Heidelberg, Germany) (Nevers et al., 2018); and Huh7.5 cells. The cells were maintained in Dulbecco’s modified Eagle’s medium Glutamax II (DMEM, Invitrogen, Carlsbad, California, USA) supplemented with 50 IU/mL penicillin, 100 µg/mL streptomycin (Invitrogen), 0.1 µg/mL fungizone and 10% fetal calf serum (FCS, Hyclone, Logan, Utah, USA) at 37°C and 5% CO2. The Huh7.5-SGR cell line was grown in the presence of 600 µg/mL geneticin (G418, Invitrogen).

### HCV infections and lysate preparations

Huh7.5 cells were infected at 37°C with the modified genotype 2a JFH1 infectious HCV strain Jad (Boukadida et al., 2014) at indicated multiplicities of infection (MOIs). Infected cells were harvested at various time points post-infection. Cells were washed with Phosphate Buffer Saline (PBS, Eurobio, Courtaboeuf, France) and lysed in cold lysis buffer (1% NP40, 10% glycerol, 100 mM Tris pH 8, 100 mM KCl, protease and phosphatase inhibitors) or RLT buffer supplemented with “- mercaptoethanol (RNeasy mini kit, Qiagen, Hilden, Germany) for western blot analysis and total RNA extraction, respectively.

### RNA extraction and quantification

Total RNA was prepared using the RNeasy mini kit (Qiagen). RNA amounts were determined by absorption spectroscopy using NanoDrop (NanoDrop Products, Wilmington, Delaware, USA). Complementary DNA was synthesized using a High Capacity cDNA Reverse Transcription kit (Applied Biosystems, Foster City, California, USA). Quantitative PCR was performed with an Applied Biosystems 7300 Thermal Cycler using TaqMan reagents (Life Technologies, Carlsbad, California, USA). HCV RNA levels were quantified by means of a quantitative real-time PCR assay using the TaqMan technology with HCV 5’-UTR specific primers (sense: 5!- GCA-GAA-AGC-GTC-TAG-CCA-TGG-CGT-3!; antisense: 5!-CTC-GCA-AGC-ACC- CTA-TCA-GGC-AGT-3!) and probe (5!-6-FAM-CAT-AGT-GGT-CTG-CGG-AAC- CGG-TGA-GT-TAMRA-3!).

Pri-miRNA levels were measured by means of the Taqman Pri-microRNA assay (Life Technologies). Results were normalized to glyceraldehyde-3-phosphate dehydrogenase (GAPDH) mRNA or 18S rRNA, as indicated.

Specific miRs were reverse-transcribed using the Taqman MicroRNA reverse transcription kit (Applied Biosystems), according to the manufacturer’s instructions. The resulting cDNAs were quantified by means of Taqman MicroRNA Assays (Life Technologies), according to the supplier’s protocol. A dilution series of synthetic oligonucleotides, ranging from 20 nM to 20 fM, was used to establish a standard curve. C_T_ values were measured for each miRNA and the standard curve per 15 pg of total RNA (it was assumed that human cell lines typically contain less than 15 pg total RNA per cell, hence the results are rough estimates of the absolute miRNA copy number per cell). For relative quantification, the results were normalized to RNU6B (U6).

HCV RNA, pri-miRNA and miRNA relative expression levels were all calculated by the !!C_T_ method. Each data point represents the average of 3 replicates in cell culture.

### siRNA and mimic-miRNA transfection

Human Custom ON-TARGET plus siKSRP or mimic miRs were resuspended in siRNA dilution buffer (Dharmacon, GE Healthcare, Chalfont-St-Giles, United Kingdom) to a 20 µM stock solution. Three additional siRNAs targeting the 3’-UTR of KSRP mRNA (siKSRP^e^) were synthesized (Dharmacon, GE Healthcare) and pooled to a 20 µM stock solution. For siRNA and mimic miRNA transfection, cells were seeded in 12-well plates at 180,000 cells/well in the presence of Dharmafect IV reagent (Dharmacon, GE Healthcare) and 12.5-25 nM of siRNA duplexes or 100 nM mimic miR, respectively, as recommended by the manufacturer. The efficiency of siRNA depletion or mimic miRNA supplementation was assessed by means of RT-qPCR or western blotting.

### Plasmid transfections and co-transfections with siRNAs

A human KSRP cDNA sequence (isoform 1), cloned into a pCMV expression plasmid and fused with an HA-tag (pKSRP-HA) was used to obtain KSRP mutants. The Quick Change XL Site-directed mutagenesis kit (Stratagene, San Diego, California, USA) was used to generate the phosphomimetic KSRP-S193D-HA and non-phosphorylable KSRP-S193A-HA proteins. For transfection, cells were seeded in 12-well plates at 180,000 cells/well and maintained in culture medium for 24 h. Cells were then washed with PBS and placed in contact with duplexes of 1 µg of plasmid and Lipofectamine 3000 (Life Technologies), according to the manufacturer’s instructions.

### Western Blotting

In total, 5-10 µg of proteins were resolved on 4–12% SDS-polyacrylamide gel and transferred to nitrocellulose membranes (GE Healthcare). Membranes were probed with various antibodies prior to development with ECL prime (GE Healthcare), including: mouse antibody against !-actin (clone AC-40, Sigma-Aldrich, St Louis, Missouri, USA); rabbit antibody against Akt (pan) (C67E7, #4691, Cell Signaling Technology, Danvers, Massachusetts, USA); rabbit antibody against phospho-Akt (Ser473) (#9271, Cell Signaling Technology); rabbit antibody against KSRP (KSRP^rabbit^, NBP1-18910, Novus Biologicals, Littleton, Colorado, USA); mouse antibody against NS5A HCV protein (AB 13833, Abcam, Cambridge, United Kingdom); rabbit antibody against HA-tag (#H6908, Sigma-Aldrich); rabbit horseradish-peroxidase conjugated antibody against GAPDH (#3683, Cell Signaling Technology); horseradish-peroxidase conjugated secondary antibodies (#31460 and #31430, Life Technologies) or rat horseradish peroxydase-conjugated antibody against HA-Tag (Clone 3F10, Sigma-Aldrich). Quantifications were performed by means of the luminescence image analyzer Image Quant Las 4000 mini and Image Quant software (GE Healthcare).

### Ribonucleoprotein complex immunoprecipitation

Ribonucleoprotein complex immunoprecipitation (RIP) was performed using the EZ-Magna RIP RNA-binding protein Immunoprecipitation Kit (Merck Millipore Billerica, Massachusetts, USA). Huh7.5-SGR cells were seeded in Petri dishes at 10^6^ cells/dish for 24 h prior to transfection with pKSRP-HA plasmids. Cells were homogenized in RIP lysis buffer 24 h post-transfection. Half of the lysate was used for immunoprecipitation with anti-KSRP^rabbit^ antibody (NBP1-18910, Novus Biologicals), anti-HA antibody (H1847-53C, US Biological, Salem, Massachusetts, USA) or anti-DROSHA antibody (AB 12286, Abcam); the other half was used with control rabbit or mouse IgG (5 µg). Immunoprecipitation was performed as recommended by the manufacturer. Immunoprecipitated RNA was extracted and purified, and the quality and amount of total immunoprecipitated RNA was determined by absorption spectroscopy using NanoDrop (NanoDrop Products). An equal amount of RNA was subjected to reverse transcription using random primers to generate cDNA libraries used for quantitative PCR reactions.

### *In situ* proximity ligation assay

*In situ* proximity ligation assay (PLA) was used to visualize protein-protein interactions and post-translational protein modifications in fixed cultured cells using secondary antibodies with attached oligonucleotides. KSRP post-translational modifications were monitored in Huh7.5-SGR and Huh7.5 cells grown on Nunc Lab-Tek Chamber Slide™ system 8 wells (Permanox, Sigma-Aldrich), 10,000 cells/well. After incubation and infection with Jad, Huh7.5 cells were fixed with 2% paraformaldehyde in PBS for 10 min at room temperature and permeabilized with cold methanol for 10 min at −20°C. Slides were blocked for 1 h with 1% BSA (fraction V), 0.3% Triton X114 in PBS at 37°C in a humidity chamber, incubated with primary antibodies directed against KSRP^rabbit^ (NBP1-18910, Novus Biologicals), KSRP^mouse^ (ab56438, Abcam), RXXS*/T* motif (110B7E, Cell Signaling Technology) or HA-tag (H1847-53C, US Biological) for 1 h at 37°C in a humidity chamber, and then processed using Duolink *in situ* Detection reagent Kit (Olink Bioscience, Uppsala, Sweden), as recommended by the manufacturer. Slides were counterstained with DAPI before mounting. Immunofluorescence experiments were performed using sheep antibody against NS5A (gift from Mark Harris, University of Leeds, United Kingdom) and Alexa Fluor 488 donkey secondary antibody against sheep (#A11015, Life Technologies). Images of *in situ* PLA and immunofluorescence experiments were taken with a Zeiss Axioskop 40 fluorescence microscope using an AxioCam MRc 3 CCD sensor and 40x or 100x objectives with filters for DAPI, FITC and Cy3.

### Primary human hepatocytes

Primary human hepatocytes (PHH, Biopredic, Rennes, France) were isolated from normal-appearing liver tissues obtained from adult patients undergoing partial hepatectomy for the treatment of metastases who were seronegative for HCV, hepatitis B virus and human immunodeficiency virus. Freshly isolated PHHs were seeded at high density on collagen-coated plates and maintained in primary culture as described previously (Podevin et al., 2010). A high-titer HCV JFH1 stock was used to infect PHHs 4 days post-seeding at an MOI of 2 focus-forming units per cell. To assess the effect of Akt inhibition, PHHs were treated with 4 !M Akt inhibitor VIII (#124018 Calbiochem, Darmstadt, Germany) or DMSO as carrier control for 24 h before HCV inoculation, and maintained in the presence of Akt inhibitor for 3 additional days before cell lysis. The inhibition of Akt activity was confirmed by western blot analysis with rabbit antibody against phospho-S473-Akt (#9271, Cell Signaling Technology), rabbit antibody against Akt (pan) (C67E7, #4691, Cell Signaling Technology, Danvers, Massachusetts, USA) and rabbit horseradish-peroxidase conjugated antibody against GAPDH (#3683, Cell Signaling Technology). HCV replication and miR-122 maturation were measured by means of RT-qPCR as previously described (Carriere et al., 2007). For KSRP silencing, siRNAs (ON-TARGETplus smartpool, Dharmacon, GE Healthcare) were diluted into PBS to a final concentration of 2 nM and mixed with 2 !l of RiboCellin siRNA Delivery Reagent (BioCellChallenge), as recommended by the manufacturer. Data are presented as means of 3 replicates in cell culture.

### Statistical analyses

The results are expressed as mean±SEM of three replicate experiments. Comparisons were made by means of the Student’s t test wherever appropriate. A p value <0.05 was considered significant. * differs from no treatment or zero time, p<0.05, ** differs from no treatment or zero time, p<0.01, *** differs from no treatment or zero time, p=<0.001.

## Acknowledgments

The authors are grateful to Dr Ralf Bartenschlager (University of Heidelberg, Germany) for kindly providing the Huh7.5 clone harboring the HCV genotype 1b bicistronic subgenomic replicon I389-neo/NS3-3’/5.1, and to Dr Annette Martin (Institut Pasteur, Paris, France) for providing the modified JFH1 (Jad) HCV strain.

---

**Figure S1:**
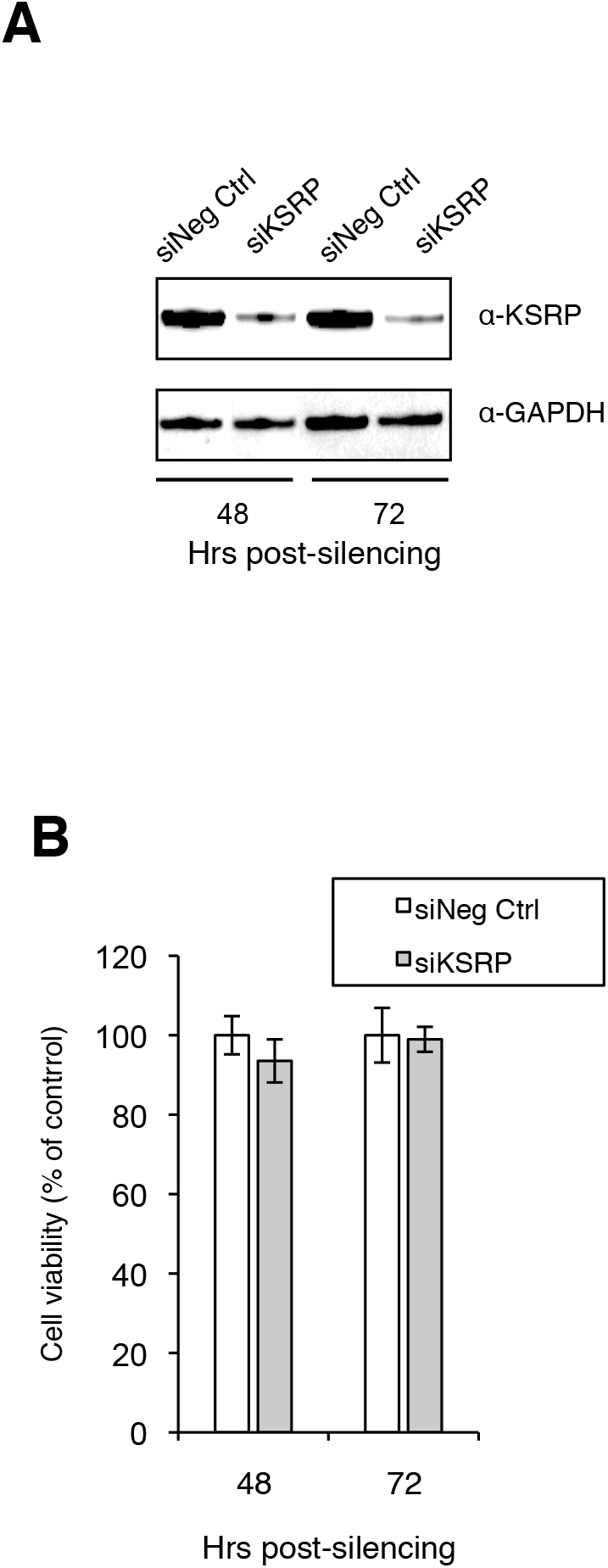
Control of KSRP silencing and cytotoxicity in Huh7.5 cells. (A) KSRP expression was quantified by immunoblot analysis using anti-KSRP and anti-GAPDH (loading control) antibodies at 48 h and 72 h post-siKSRP transfection. (B) Cytotoxicity assay.

**Supplementary Figure S2:**
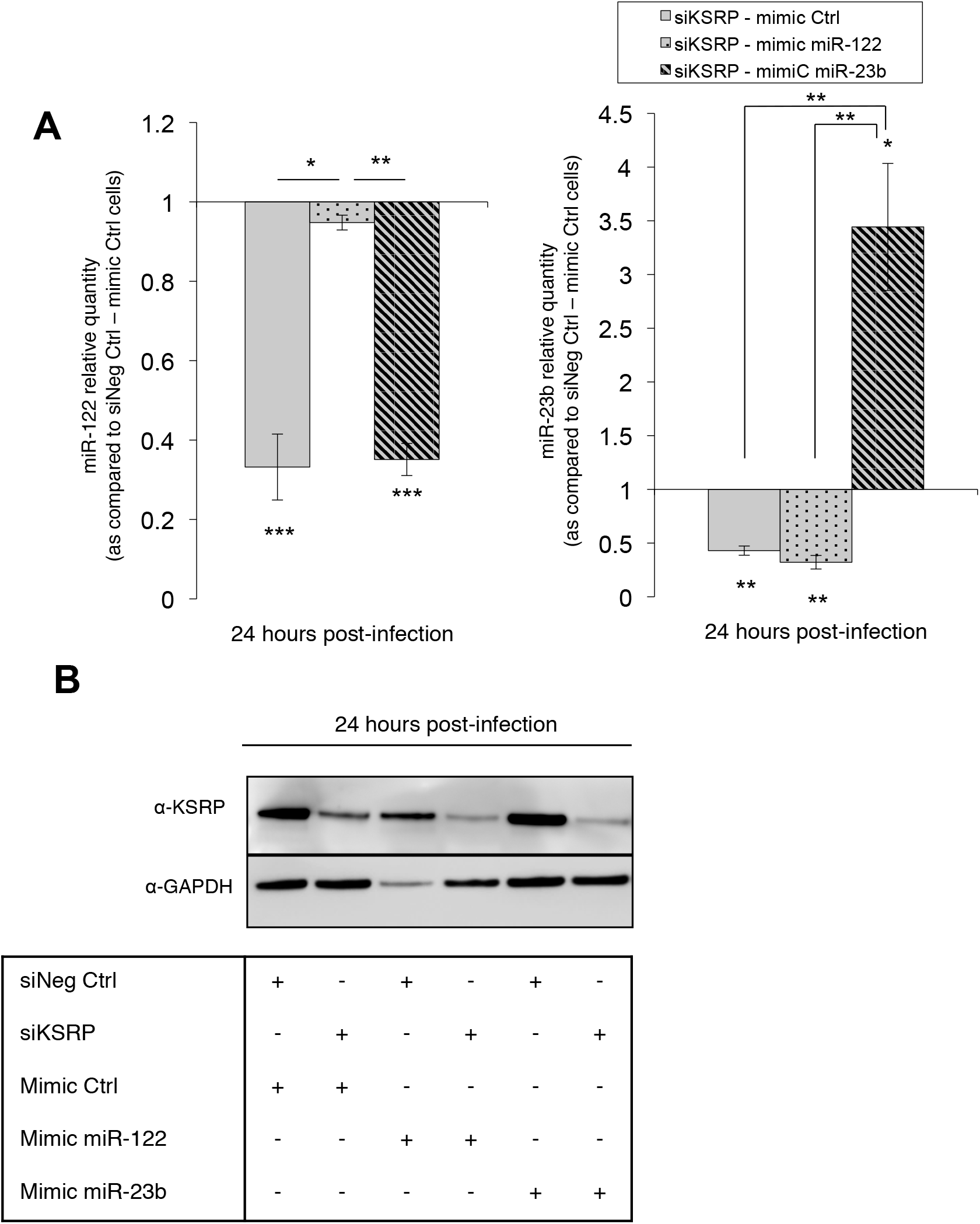
Control of mimic miRNA transfection and KSRP silencing in HCV Jad-infected Huh7.5 cells. (A) Mimic miR-122 and miR-23b transfections were quantified by means of RT-qPCR. miRNA quantities were normalized to RNU6B (U6) and are presented as relative expression to siNeg Ctrl-mimic Ctrl. (B) Immunoblot analysis of KSRP silencing in Huh7.5 cells co-transfected with mimic miRNA. KSRP was quantified 24 h post-infection, i.e. 48 h post-co-transfection of siRNA and mimic miRNA, using anti-KSRP and anti-GAPDH antibodies. *: p<0.05; **: p<0.01; ***: p<0.001 for comparisons or significance of fold-increases.

**Supplementary Figure S3:**
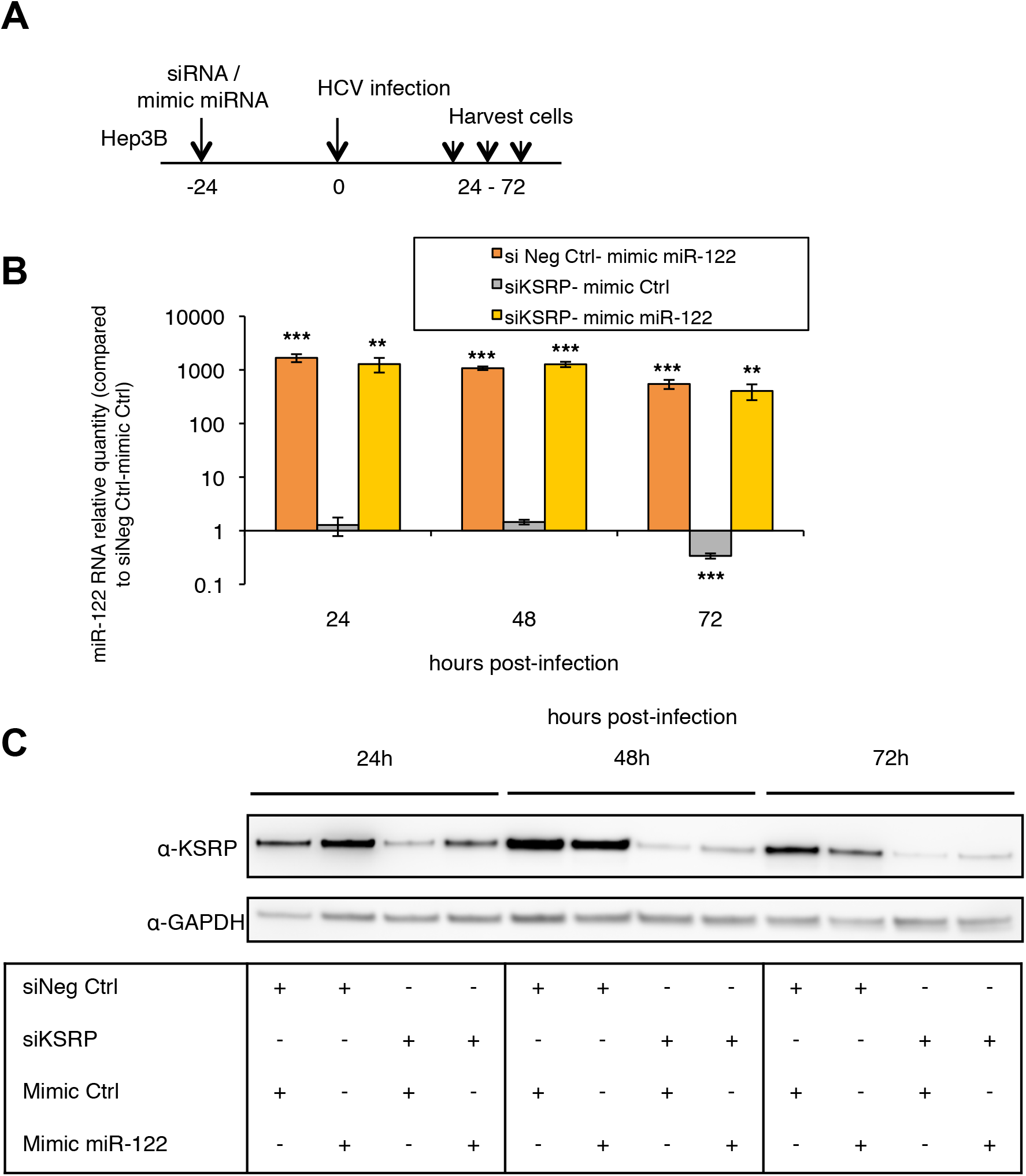
Control of mimic miR-122 transfection and KSRP silencing in HCV Jad-infected Hep3B cells. (A) Experimental design: Hep3b cells were co-transfected with scrambled control (siNeg Ctrl), siKSRP, mimic Ctrl or mimic miR-122 and infected 24 h later with the Jad HCV strain at an MOI of 0.1. (B) Mimic miR-122 transfection was quantified by means of RT-qPCR. miRNA quantities were normalized to RNU6B (U6) and are presented as relative expression to siNeg Ctrl - mimic Ctrl at each time point. (C) Immunoblot analysis of KSRP silencing in cells co-transfected with mimic miRNA. KSRP was quantified 24 to 72 h post-infection, i.e. 48 to 96 h post-co-transfection of siRNA and mimic miRNA, using anti-KSRP and anti-GAPDH antibodies. *: p<0.05; **: p<0.01; ***: p<0.001 for significance of fold-increases.

**Supplementary Figure S4:**
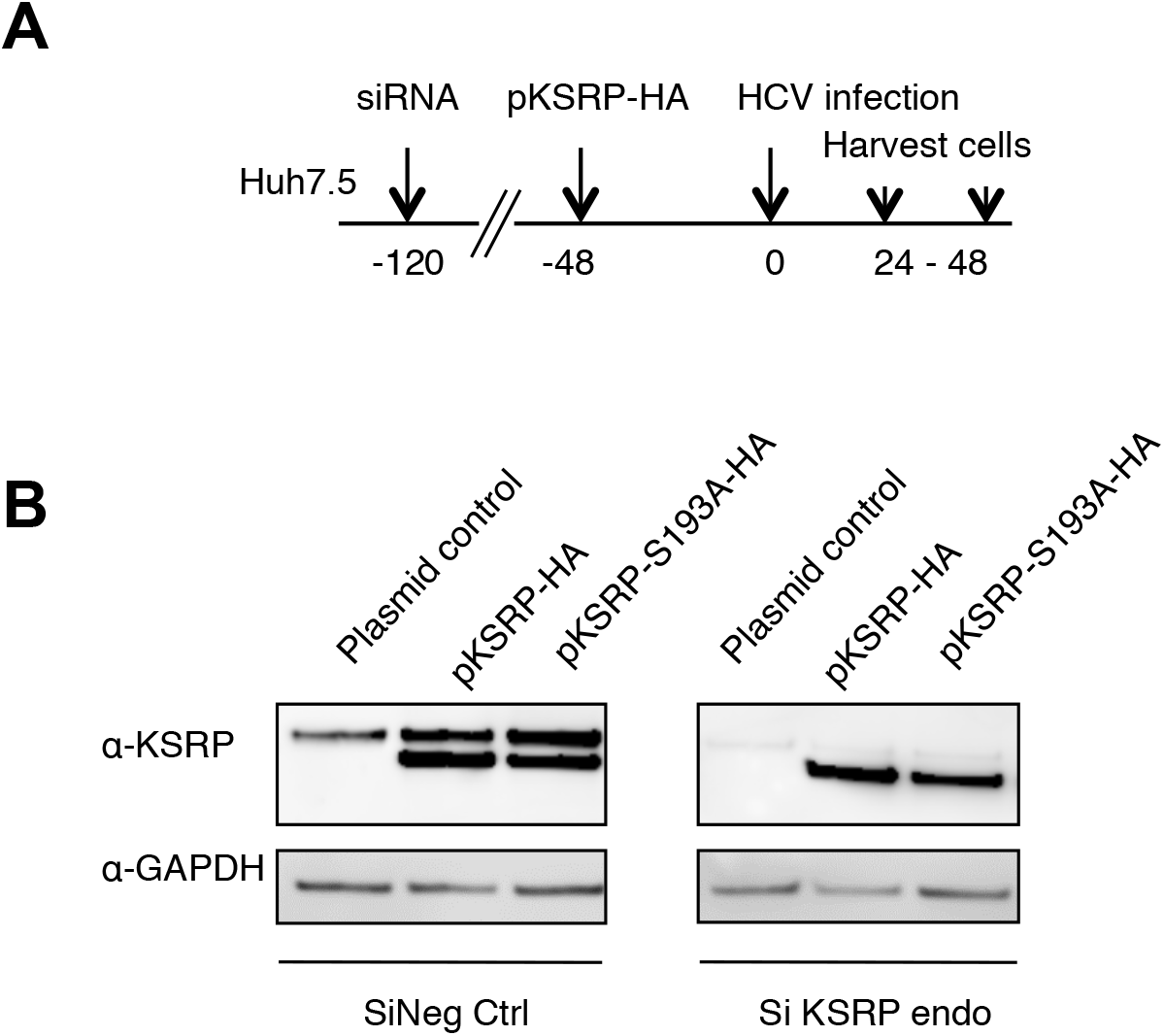
Control of KSRP silencing and transfection of plasmids expressing KSRP-HA variants in Huh7.5 cells 48 h post-infection with the HCV Jad strain at an MOI of 0.1. (A) Experimental design: Huh7.5 cells were transfected with scrambled control (siNeg Ctrl) or siKSRP^e^ that targets the 3’UTR of KSRP mRNA, for 72 h. The control plasmid and plasmids expressing KSRP-HA or KSRP-S193A proteins were transfected for 48 h before infection with the HCV Jad strain at an MOI of 0.1. (B) KSRP and KSRP-HA proteins were quantified 48 h post-infection, i.e. 168 h post-transfection of siNeg Ctrl or siKSRP^e^, using anti-KSRP and anti-GAPDH antibodies.

**Supplementary Figure S5:**
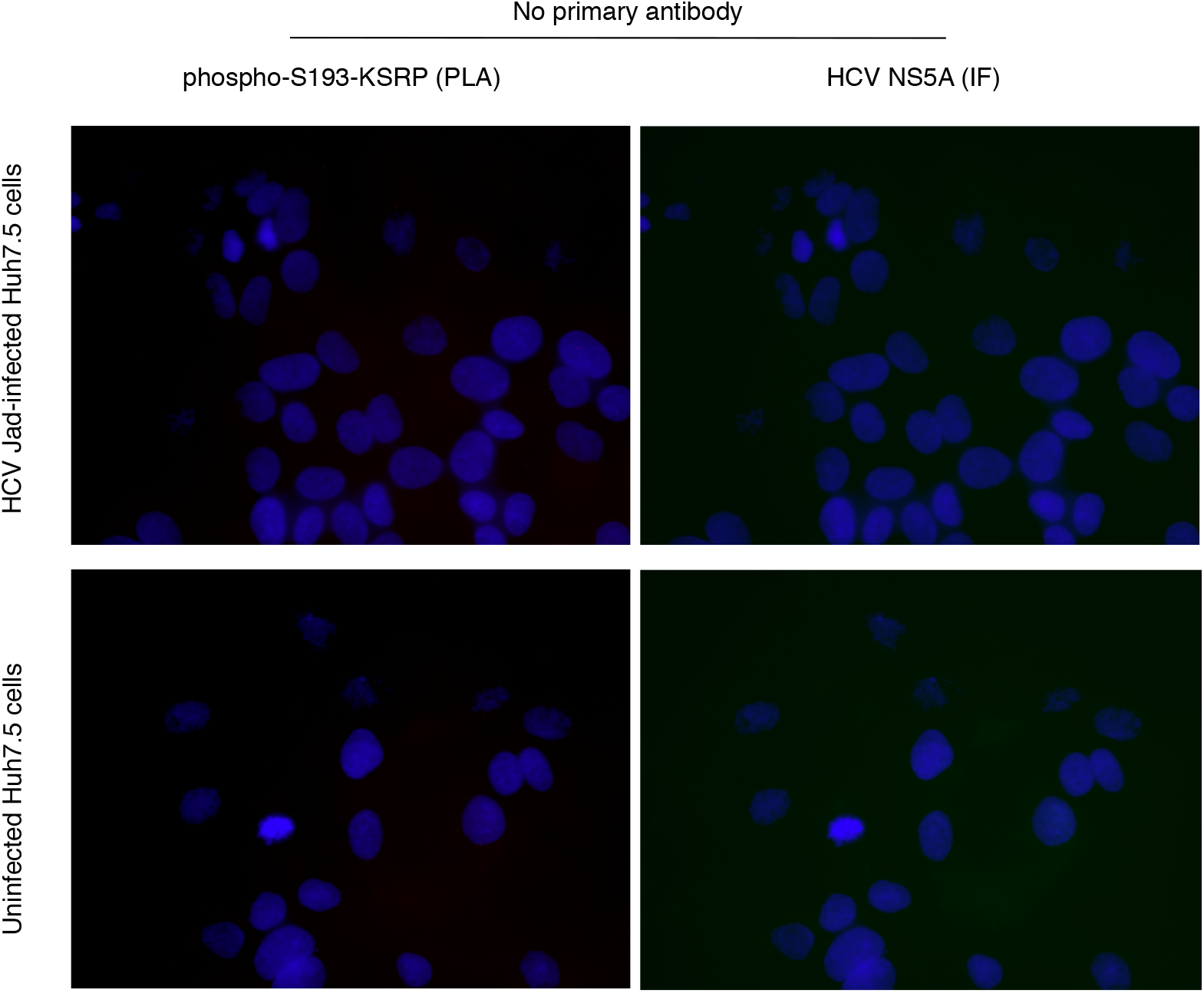
Effect of HCV infection on the amount of phospho-S193-KSRP in hepatoma cell line, as assessed by *in situ* PLA. Control representative *in situ* PLA images showing phospho-S193-KSRP in Huh7.5 cells infected with the HCV genotype 2a Jad strain at an MOI of 0.1, as compared to uninfected cells. Cells were probed without antibodies directed against the HCV NS5A protein (immunofluorescence, IF signal, green) or phospho-S193-KSRP (KSRP^mouse^ x RXXS*/ T*) (PLA signal, red). Cell nuclei were stained with DAPI (blue).

## References

Amador-Canizares, Y., Bernier, A., Wilson, J. A., & Sagan, S. M. (2018). miR-122 does not impact recognition of the HCV genome by innate sensors of RNA but rather protects the 5’ end from the cellular pyrophosphatases, DOM3Z and DUSP11. Nucleic Acids Res. doi: 10.1093/nar/gky273

Bala, S., Marcos, M., & Szabo, G. (2009). Emerging role of microRNAs in liver diseases. World J Gastroenterol, 15(45), 5633–5640. doi 10.3748/wjg.15.5633

Bose, S. K., Meyer, K., Di Bisceglie, A. M., Ray, R. B., & Ray, R. (2012). Hepatitis C virus induces epithelial-mesenchymal transition in primary human hepatocytes. J Virol, 86(24), 13621–13628. doi 10.1128/JVI.02016-12

Boukadida, C., Marnata, C., Montserret, R., Cohen, L., Blumen, B., Gouttenoire, J., …Martin, A. (2014). NS2 proteins of GB virus B and hepatitis C virus share common protease activities and membrane topologies. J Virol, 88(13), 7426–7444. doi 10.1128/JVI.00656-14

Briata, P., Chen, C. Y., Ramos, A., & Gherzi, R. (2013). Functional and molecular insights into KSRP function in mRNA decay. Biochim Biophys Acta, 1829(6-7), 689–694. doi: 10.1016/j.bbagrm.2012.11.003

Briata, P., Forcales, S. V., Ponassi, M., Corte, G., Chen, C. Y., Karin, M., … Gherzi, R. (2005). p38-dependent phosphorylation of the mRNA decay-promoting factor KSRP controls the stability of select myogenic transcripts. Mol Cell, 20(6), 891–903. doi 10.1016/j.molcel.2005.10.021

Briata, P., Lin, W. J., Giovarelli, M., Pasero, M., Chou, C. F., Trabucchi, M., … Gherzi, R. (2012). PI3K/AKT signaling determines a dynamic switch between distinct KSRP functions favoring skeletal myogenesis. Cell Death Differ, 19(3), 478–487. doi 10.1038/cdd.2011.117

Carriere, M., Pene, V., Breiman, A., Conti, F., Chouzenoux, S., Meurs, E., … Podevin, P. (2007). A novel, sensitive, and specific RT-PCR technique for quantitation of hepatitis C virus replication. J Med Virol, 79(2), 155–160. doi 10.1002/jmv.20773

Cheng, D., Zhang, L., Yang, G., Zhao, L., Peng, F., Tian, Y., … Gong, G. (2015). Hepatitis C virus NS5A drives a PTEN-PI3K/Akt feedback loop to support cell survival. Liver Int, 35(6), 1682–1691. doi 10.1111/liv.12733

Colman, H., Le Berre-Scoul, C., Hernandez, C., Pierredon, S., Bihouee, A., Houlgatte, R., … Feray, C. (2013). Genome-wide analysis of host mRNA translation during hepatitis C virus infection. J Virol, 87(12), 6668–6677. doi 10.1128/JVI.00538-13

Frentzen, A., Anggakusuma Gurlevik, E., Hueging, K., Knocke, S., Ginkel, C., … Pietschmann, T. (2014). Cell entry, efficient RNA replication, and production of infectious hepatitis C virus progeny in mouse liver-derived cells. Hepatology, 59(1), 78–88. doi 10.1002/hep.26626

Friebe, P., Lohmann, V., Krieger, N., & Bartenschlager, R. (2001). Sequences in the 5’ nontranslated region of hepatitis C virus required for RNA replication. J Virol, 75(24), 12047–12057. doi 10.1128/JVI.75.24.12047-12057.2001

Fukuhara, T., Kambara, H., Shiokawa, M., Ono, C., Katoh, H., Morita, E., … Matsuura, Y. (2012). Expression of microRNA miR-122 facilitates an efficient replication in nonhepatic cells upon infection with hepatitis C virus. J Virol, 86(15), 7918–7933. doi 10.1128/JVI.00567-12

Gherzi, R., Chen, C. Y., Trabucchi, M., Ramos, A., & Briata, P. (2010). The role of KSRP in mRNA decay and microRNA precursor maturation. Wiley Interdiscip Rev RNA, 1(2), 230–239. doi 10.1002/wrna.2

Han, Y., Niu, J., Wang, D., & Li, Y. (2016). Hepatitis C Virus Protein Interaction Network Analysis Based on Hepatocellular Carcinoma. PLoS One, 11(4), e0153882. doi 10.1371/journal.pone.0153882

He, Y., Nakao, H., Tan, S. L., Polyak, S. J., Neddermann, P., Vijaysri, S., … Katze M. G. (2002). Subversion of cell signaling pathways by hepatitis C virus nonstructural 5A protein via interaction with Grb2 and P85 phosphatidylinositol 3-kinase. J Virol, 76(18), 9207–9217.

Higgs, M. R., Lerat, H., & Pawlotsky, J. M. (2013). Hepatitis C virus-induced activation of beta-catenin promotes c-Myc expression and a cascade of pro-carcinogenetic events. Oncogene, 32(39), 4683–4693. doi 10.1038/onc.2012.484

Honda, M., Beard, M. R., Ping, L. H., & Lemon, S. M. (1999). A phylogenetically conserved stem-loop structure at the 5’ border of the internal ribosome entry site of hepatitis C virus is required for cap-independent viral translation. J Virol, 73(2), 1165–1174.

Hoofnagle, J. H. (2002). Course and outcome of hepatitis C. Hepatology, 36(5 Suppl 1), S21-29. doi: 10.1053/jhep.2002.36227

Jopling, C. L., Schutz, S., & Sarnow, P. (2008). Position-dependent function for a tandem microRNA miR-122-binding site located in the hepatitis C virus RNA genome. Cell Host Microbe, 4(1), 77–85. doi 10.1016/j.chom.2008.05.013

Jopling, C. L., Yi, M., Lancaster, A. M., Lemon, S. M., & Sarnow, P. (2005). Modulation of hepatitis C virus RNA abundance by a liver-specific MicroRNA. Science, 309(5740), 1577–1581. doi 10.1126/science.1113329

Kim, G. W., Lee, S. H., Cho, H., Kim, M., Shin, E. C., & Oh, J. W. (2016). Hepatitis C Virus Core Protein Promotes miR-122 Destabilization by Inhibiting GLD-2. PLoS Pathog, 12(7), e1005714. doi 10.1371/journal.ppat.1005714

Kroll, T. T., Zhao, W. M., Jiang, C., & Huber, P. W. (2002). A homolog of FBP2/KSRP binds to localized mRNAs in Xenopus oocytes. Development, 129(24), 5609–5619.

Li, Y., Masaki, T., & Lemon, S. M. (2013). miR-122 and the Hepatitis C RNA genome: more than just stability. RNA Biol, 10(6), 919–923. doi 10.4161/rna.25137

Li, Y., Yamane, D., Masaki, T., & Lemon, S. M. (2015). The yin and yang of hepatitis C: synthesis and decay of hepatitis C virus RNA. Nat Rev Microbiol, 13(9), 544–558. doi 10.1038/nrmicro3506

Liu, Y., & Liu, Q. (2011). ATM signals miRNA biogenesis through KSRP. Mol Cell, 41(4), 367–368. doi 10.1016/j.molcel.2011.01.027

Liu, Z., Tian, Y., Machida, K., Lai, M. M., Luo, G., Foung, S. K., & Ou, J. H. (2012). Transient activation of the PI3K-AKT pathway by hepatitis C virus to enhance viral entry. J Biol Chem, 287(50), 41922–41930. doi 10.1074/jbc.M112.414789

Luna, J. M., Scheel, T. K., Danino, T., Shaw, K. S., Mele, A., Fak, J. J., … Darnell, R. B. (2015). Hepatitis C virus RNA functionally sequesters miR-122. Cell, 160(6), 1099–1110. doi 10.1016/j.cell.2015.02.025

Marquez, R. T., Bandyopadhyay, S., Wendlandt, E. B., Keck, K., Hoffer, B. A., Icardi, M. S., … McCaffrey, A. P.. (2010). Correlation between microRNA expression levels and clinical parameters associated with chronic hepatitis C viral infection in humans. Lab Invest, 90(12), 1727–1736. doi 10.1038/labinvest.2010.126

Masaki, T., Arend, K. C., Li, Y., Yamane, D., McGivern, D. R., Kato, T., … Lemon, S. M. (2015). miR-122 stimulates hepatitis C virus RNA synthesis by altering the balance of viral RNAs engaged in replication versus translation. Cell Host Microbe, 17(2), 217–228. doi 10.1016/j.chom.2014.12.014

Milward, A., Mankouri, J., & Harris, M. (2010). Hepatitis C virus NS5A protein interacts with beta-catenin and stimulates its transcriptional activity in a phosphoinositide-3 kinase-dependent fashion. J Gen Virol, 91(Pt 2), 373–381. doi 10.1099/vir.0.015305-0

Min, H., Turck, C. W., Nikolic, J. M., & Black, D. L. (1997). A new regulatory protein, KSRP, mediates exon inclusion through an intronic splicing enhancer. Genes Dev, 11(8), 1023–1036.

Nevers, Q., Ruiz, I., Ahnou, N., Donati, F., Brillet, R., Softic, L., … Ahmed-Belkacem, A. (2018). Characterization of the Anti-Hepatitis C Virus Activity of New Nonpeptidic Small-Molecule Cyclophilin Inhibitors with the Potential for Broad Anti-Flaviviridae Activity. Antimicrob Agents Chemother, 62(7). doi 10.1128/AAC.00126-18

Podevin, P., Carpentier, A., Pene, V., Aoudjehane, L., Carriere, M., Zaidi, S., … Calmus, Y. (2010). Production of infectious hepatitis C virus in primary cultures of human adult hepatocytes. Gastroenterology, 139(4), 1355–1364. doi 10.1053/j.gastro.2010.06.058

Repetto, E., Briata, P., Kuziner, N., Harfe, B. D., McManus, M. T., Gherzi, R., … Trabucchi, M. (2012). Let-7b/c enhance the stability of a tissue-specific mRNA during mammalian organogenesis as part of a feedback loop involving KSRP. PLoS Genet, 8(7), e1002823. doi 10.1371/journal.pgen.1002823

Sedano, C. D., & Sarnow, P. (2014). Hepatitis C virus subverts liver-specific miR-122 to protect the viral genome from exoribonuclease Xrn2. Cell Host Microbe, 16(2), 257–264. doi 10.1016/j.chom.2014.07.006

Shi, Q., Hoffman, B., & Liu, Q. (2016). PI3K-Akt signaling pathway upregulates hepatitis C virus RNA translation through the activation of SREBPs. Virology, 490, 99–108. doi 10.1016/j.virol.2016.01.012

Shimakami, T., Yamane, D., Jangra, R. K., Kempf, B. J., Spaniel, C., Barton, D. J., & Lemon, S. M. (2012). Stabilization of hepatitis C virus RNA by an Ago2-miR-122 complex. Proc Natl Acad Sci U S A, 109(3), 941–946. doi 10.1073/pnas.1112263109

Street, A., Macdonald, A., Crowder, K., & Harris, M. (2004). The Hepatitis C virus NS5A protein activates a phosphoinositide 3-kinase-dependent survival signaling cascade. J Biol Chem, 279(13), 12232–12241. doi 10.1074/jbc.M312245200

Thibault, P. A., Huys, A., Dhillon, P., & Wilson, J. A. (2013). MicroRNA-122-dependent and - independent replication of Hepatitis C Virus in Hep3B human hepatoma cells. Virology, 436(1), 179–190. doi 10.1016/j.virol.2012.11.007

Trabucchi, M., Briata, P., Garcia-Mayoral, M., Haase, A. D., Filipowicz, W., Ramos, A., … Rosenfeld, M. G. (2009). The RNA-binding protein KSRP promotes the biogenesis of a subset of microRNAs. Nature, 459(7249), 1010–1014. doi 10.1038/nature08025

van der Ree, M. H., de Vree, J. M., Stelma, F., Willemse, S., van der Valk, M., Rietdijk, S., … Reesink, H. W. (2017). Safety, tolerability, and antiviral effect of RG-101 in patients with chronic hepatitis C: a phase 1B, double-blind, randomised controlled trial. Lancet, 389(10070), 709–717. doi 10.1016/S0140-6736(16)31715-9

Varnholt, H., Drebber, U., Schulze, F., Wedemeyer, I., Schirmacher, P., Dienes, H. P., & Odenthal, M. (2008). MicroRNA gene expression profile of hepatitis C virus-associated hepatocellular carcinoma. Hepatology, 47(4), 1223–1232. doi 10.1002/hep.22158

Zhang, X., Wan, G., Berger, F. G., He, X., & Lu, X. (2011). The ATM kinase induces microRNA biogenesis in the DNA damage response. Mol Cell, 41(4), 371–383. doi 10.1016/j.molcel.2011.01.020

